# KaroSpace: a rapid-access framework for interactive exploration of multi-sample spatial omics data

**DOI:** 10.64898/2026.03.10.710790

**Authors:** Christoffer M. Langseth, Bastien Hervé, Hanna P. Piechaczyk, Yuk Kit Lor, Ting Sun, Gonçalo Castelo-Branco

## Abstract

Spatial omics technologies enable high-resolution mapping of molecular and cellular organization within tissues, yet interactive exploration of these data remains challenging due to computational bottlenecks, and reliance on proprietary software infrastructures. We present KaroSpace, a framework for cell-centric exploration of multi-sample and multi-modality spatial omics data, agnostic to upstream analysis pipelines and operating on preprocessed spatial features. By combining a lightweight design with flexible deployment options, KaroSpace allows immediate interactive spatial data exploration, straightforward implementation to the broader scientific community, supports transparent data sharing, and complements existing computational workflows for hypothesis generation and collaborative analysis.

## Main

Recent advances in spatial omics technologies have enabled the measurement of gene expression and other molecular features within their native tissue context at cellular and subcellular resolution^1–4^. The platforms for generating spatial omics data have also matured significantly, moving from a niche technical practice in specialised laboratories, to widely adopted across diverse biological research settings, with emerging applications in translational research. Together, these developments have led to the rapid growth of increasingly large and complex spatial omics datasets that span an increasing quantity of samples, conditions, disease stages, anatomical regions, experimental models, and sample-level covariates.

Despite these advances, effective interactive exploration of spatial omics data remains a challenge. The size and complexity of the data, and reliance on proprietary or tightly coupled software infrastructures limit fast and widespread accessibility, complicate data sharing and hinder collaborative interpretation of the underlying biology. Current viewers provided with commercial spatial omics platforms offer deep single-sample data inspection. However, they typically lack native support for multi-sample and multimodality exploration. In addition, these viewers often depend on restricted access options to the complete experimental data stack. A number of visualization tools for spatial omics data have been developed, including Vitessce^5^, and TissUUmaps^6,7^, which provide feature-rich environments for spatial data inspection. CellxGene^8^ and UCSC Cell Browser^9^ are primarily embedding-centric single-cell explorers (e.g., UMAP) but have subsequently incorporated spatial functionality.

Here, we present KaroSpace, a streamlined, accessible, and portable framework built around a cell-centric design (or capture array/pixel resolution, as in Visium and DBiT-seq, respectively) across multiple spatial multiomics platforms. KaroSpace enables gene-level analysis and converts spatial omics AnnData objects into a single, self-contained HTML viewer for interactive exploration in a web browser. By eliminating reliance on a backend server, this approach enhances portability and shareability.

Reports can be generated using the KaroSpace Python API or command-line interface. To further simplify viewer creation, we developed a companion application, **KaroSpaceBuilder**, which supports macOS (ARM), Windows, and Linux and is available at https://github.com/christoffermattssonlangseth/KaroSpaceBuilder. The starting point is a.`h5ad` file (**Fig. 1a**), which can be exported from Scanpy^10^ or Seurat^11^. Once generated, the interactive report enables users to explore the dataset either through the overview panel, which displays all samples, which can then be compared, or by selecting individual samples to analyse them in more detail.

**Fig. 1.**
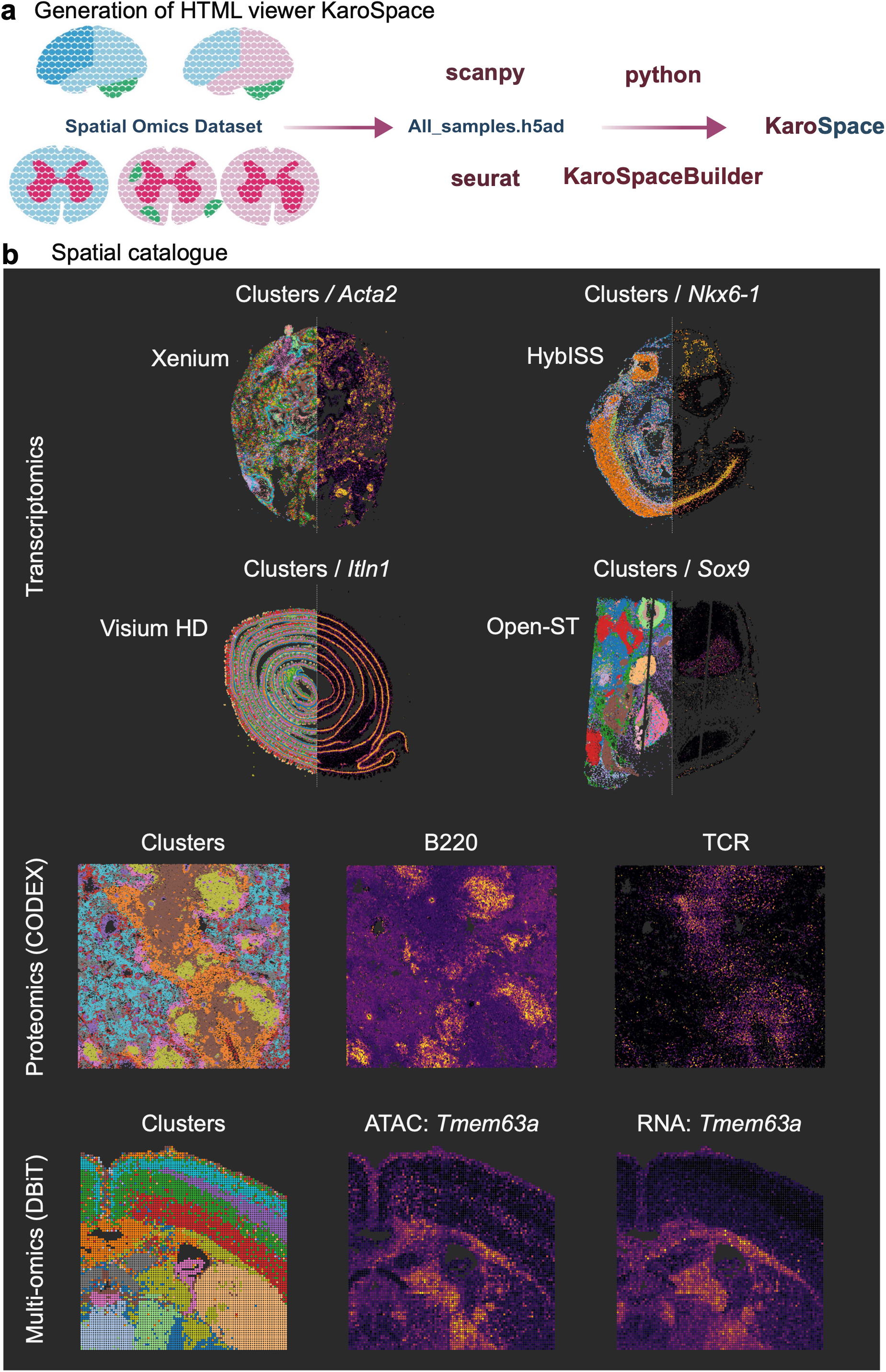
Cross-platform workflow for generating interactive spatial omics reports with KaroSpace. **a**, Spatial omics datasets from multiple platforms are processed in Scanpy or Seurat, converted from.`h5ad` objects into web-compatible formats, and compiled into interactive HTML reports using the KaroSpace Python API or KaroSpaceBuilder. **b**, Representative datasets supported by KaroSpace, including spatial transcriptomics (Xenium, HybISS, Visium HD, Open-ST), spatial proteomics (CODEX), and multiomic platforms (DBiT).

To demonstrate the flexibility of KaroSpace, we generated KaroSpace viewers across a diverse set of spatial omics datasets spanning multiple technological platforms (**Table 1; Fig 1b**). These included DBiT multiomics (transcriptome and epigenome)^12^, *In situ* sequencing–based transcriptomics datasets^13–15^, spatial proteomics data (CODEX)^16^, NanoString CosMx spatial molecular imaging datasets^17^, MERFISH data^18^, as well as publicly available datasets using the Xenium platform^19–21^. Additionally, we included Open-ST data^22^ and publicly available Visium HD data (10x Genomics website). These datasets can be viewed at KaroSpace.se.

**Table 1.** Summary of spatial omics datasets included in this study.

| Platform | Dataset (species and tissue) | Number of cells | Number of Sections | Features displayed | Report Size (MB) | Ref. |
| --- | --- | --- | --- | --- | --- | --- |
| DBiT (RNA/ATAC) | Mouse brain (Lysolecithin injection) | 50,756 | 6 | 400 (200 RNA and corresponding ATAC) | 368.1 | 12 |
| CODEX (protein) | Mouse spleen | 707,474 | 9 | 29 | 447.7 | 16 |
| MERFISH (RNA) | Mouse brain (coronal) | 6,893,712 | 218 | 20 | 496.9 | 18 |
| MERFISH (RNA) | Mouse brain (sagittal) | 2,449,745 | 28 | 50 | 262.5 | 18 |
| CosMx (RNA) | Human lung: non-small cell lung cancer | 241,008 | 2 | 50 | 32.5 | 17 |
| Xenium (RNA) | Whole mouse pup | 1,326,723 | 1 | 50 | 281.9 | NA |
| HybISS (RNA) | Whole mouse pup | 321,788 | 27 | 50 | 89.1 | 14 |
| CARTANA (RNA) | Mouse spinal cord (EAE) | 270,878 | 44 | 6 | 56.8 | 13 |
| Xenium (RNA) | Human pancreatic adenocarcinoma | 174,374 | 1 | 200 | 111.6 | NA |
| Xenium (RNA) | Human kidney | 453,447 | 1 | 20 | 48.0 | NA |
| Xenium (RNA) | Human fetal lung | 820,655 | 5 | 100 | 235.3 | 19 |
| Xenium (RNA) | Human fetal lung (acute lung injury) | 1,499,516 | 2 | 50 | 456.6 | 20 |
| Xenium (RNA) | Human lung (pulmonary fibrosis) | 88,475 | 3 | 100 | 23.8 | 21 |
| Visium HD (RNA) | Mouse intestine | 91,032 | 1 | 300 | 85 | NA |
| Visium HD (RNA) | Mouse brain | 6,484 | 2 | 300 | 25.78 | NA |
| Open-ST (RNA) | Mouse fetal brain | 84,220 | 2 | 500 | 83.2 | 22 |
| Open-ST (RNA) | Human metastatic lymph node | 1,097,769 | 19 | 25 | 333.5 | 22 |

Karospace’s report opens in the multi-sample view, providing a cohort-level overview (**Fig. 2a**). When sample-level covariates are included during report generation, users can filter samples accordingly. The **Insights** tab summarizes metadata-level statistics, including neighborhood connection proportions and enrichment scores derived from the spatial neighborhood graph, alongside gene-level views such as marker dot plots and category-wise expression exploration (**Fig. 2b**). The **Hops** feature allows one to visualize which cells are neighbouring an individual cell, at three different hop levels (**Fig. 2b**). Within the **Legend** panel, users can toggle category labels or activate the Spotlight function for rapid color reassignment. In the **Sample** view, the magic wand tool enables polygon-based cell selection with automatic aggregation of cellular composition (**Fig. 2b**). The **Annotate** function supports manual delineation of tissue regions, which can be exported as JSON files and mapped back to the original AnnData object for downstream analysis (**Fig. 2b**). For comparative visualization, a **Variable slider** allows users to sweep across the tissue while displaying two opposing variables—such as annotations versus gene expression, gene expression versus chromatin accessibility at gene promoters (gene activity), gene–gene comparisons, or alternative clustering resolutions—within the same spatial context (**Fig. 2b**).

**Fig. 2.**
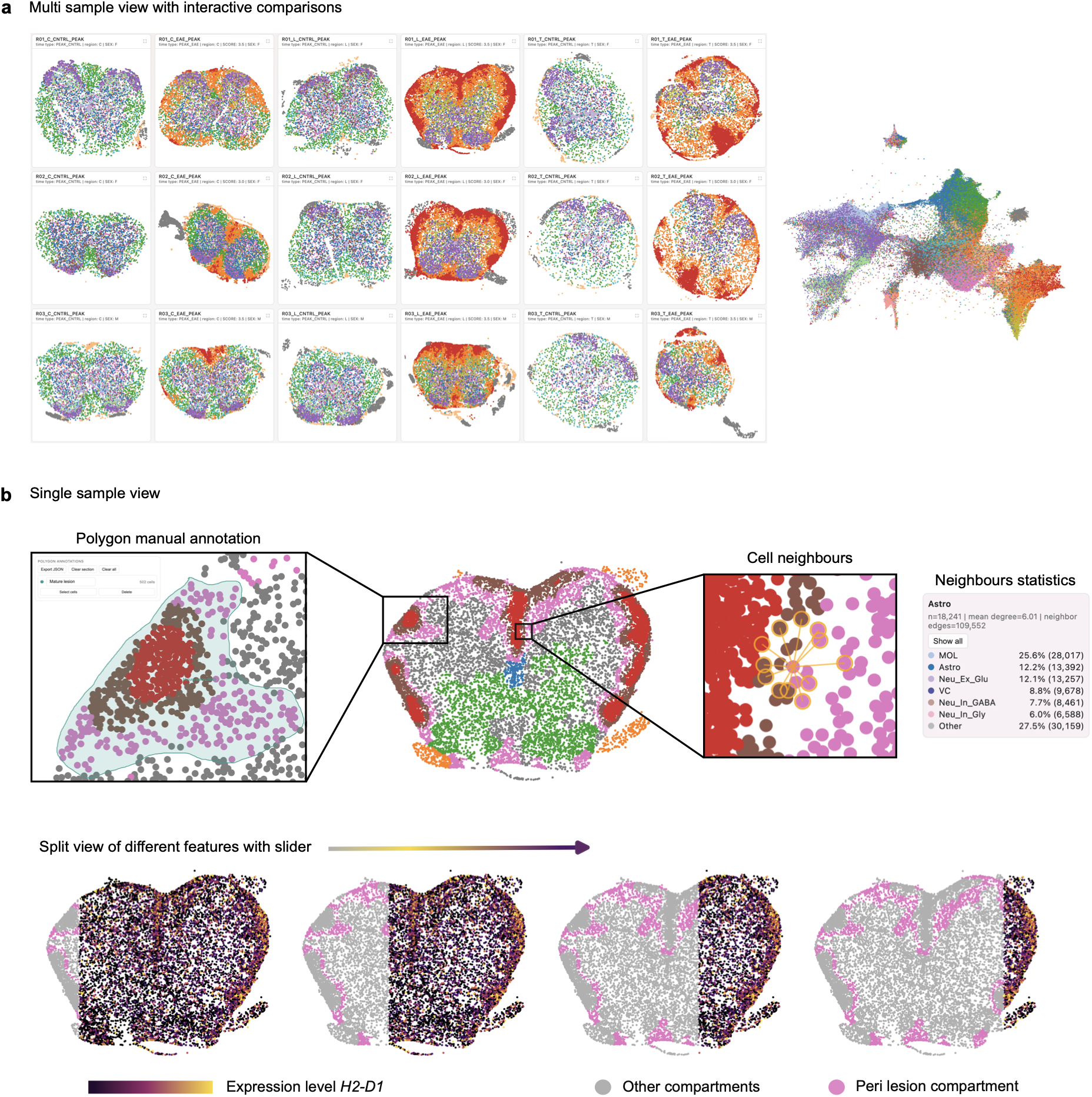
KaroSpace enables interactive multi-sample visualization, annotation, spatial neighborhood profiling, and dual-feature visualization. **a**, The multi-sample view provides a cohort-level overview of all samples and enables interactive exploration of cellular annotations and gene expression across the dataset. An accompanying embedding view displays cells in two-dimensional space. **b**, The sample view supports in-depth exploration, including polygon-based cell selection, simultaneous visualization of annotations and gene expression, manual tissue annotation with JSON export, neighborhood and metadata summaries, marker gene dot plots, and projection of embedding selections onto the spatial coordinate system.

KaroSpace offers a unique platform for immediate accessibility and ease-sharing of spatial datasets. A major feature of KaroSpace is the effective handling of datasets comprising millions of cells. Nevertheless, expanding the number of genes substantially increases file size, resulting in slower loading times and increased interface latency, and impacting on the portability, due to the size of the output files. These trade-offs reflect deliberate design choices prioritizing portability and shareability, while also highlighting areas for future optimization. Additionally, performance can vary across web browsers. For instance, while opening reports in Chrome is fast and efficient, Safari can be particularly time-consuming due to the heavy reliance on Canvas-based rendering.

In sum, as spatial omics datasets continue to scale in size and complexity, KaroSpace offers a fast, user-friendly practical solution for interactive, cross-sample visualization tailored to modern spatial biology workflows. KaroSpace will open the way to the widespread accessibility of already published and new spatial omics datasets, thereby dramatically expanding the range of users who will be able to analyse, interpret and infer biological-relevant mechanisms from spatial omics data.

## Methods

### Input data model and section construction

The input data is provided as a.h5ad file. Karospace requires 2D spatial coordinates in `adata.obsm[“spatial”]` and a tissue section identifier column. Sections are defined by a unique groupby values and optional metadata columns are attached per section for filtering and panel annotations. If there is a `adata.obsm[“X_umap”]` present in the object, a linked UMAP visualization is included as well.

### Export pipeline

For each section, KaroSpace extracts the cellular coordinates, the selected metadata, the precomputed color layers which can be categorical or continues adata.obs columns, or gene-expression vectors from `adata.var_names,` using `adata.layers[‘normalized’]` when available, otherwise `adata.X`. Optional per-section downsampling can also be applied before the export. To reduce the output size and the parse time in browsers, large numeric arrays, such as the coordinates, color values, UMAP coordinates, cell indices and graph edges are optionally serialized as a base64 encoded typed array. Furthermore, gene vectors are encoded as dense or sparse arrays and in auto mode, sparse encoding is selected for genes with a high zero fraction.

### Graph-aware analyses and marker calculations

If a neighborhood graph is available in `adata.obsp[‘spatial_connectivities’]`, KaroSpace computes neighborhood composition statistics for selected categorical groupings. Category-by-category edge counts are obtained by matrix multiplication of one-hot labelled with the graph adjacency matrix, and optional permutation testing yields z-score enrichment estimates. Optimal marker genes per category are computed using `scanpy.tl.rank_genes_groups` using a t-test.

### Interactive viewer behaviour

The exported HTML provides a linked multi-section grid and detailed modal views that allow for zooming and panning. Furthermore, color switching, gene expression rendering, lasso-based selection, category filtering, optional UMAP panel, optional graph overlays, and split view comparison of two variables, which can be cell annotation or gene expression. Polygon annotations drawn in the viewer can be exported as a JSON and then mapped back to the AnnData objects.

### Building KaroSpace viewers with KaroSpaceBuilder

In addition to the Python API that is associated with the KaroSpace repository, we also built a desktop GUI that configures and executes KaroSpace exports from AnnData inputs. Users first provide an input file and output directory, then run an inspection step that reads the `adata.obs` and `adata.var_names` that then populates the searchable selectors for grouping variables, colors and gene panels.

Coordinate handling is automatic but in an explicit way. The builder will prioritize `adata.obsm[‘spatial’]`, but if that is unavailable, it will detect the `centroid_x` and `centroid_y` from the `adata.obs`.When the centroid mode is used, the application creates a temporary.`h5ad` which contains these centroids and copies that into `obsm[‘spatial’]`.

Gene overlay selection supports four modes, highly variable gene-based, top-mean genes, file-based lists and manual lists. Duplicate genes are removed and selected genes are checked against the `adata.var_names` and invalid entries terminate export with an error. Additional analytics settings, such as marker genes, neighbourhood statistics, are sanitized before the export by removing missing columns and single-level groupings. For neighborhood statistics, estimated dense-memory cost is checked to avoid excessive allocations.

The export is executed in the background and calls `karospace.load_spatial_data()` and then `karospace.export_to_html()` with GUI defined parameters. Output is written as a timestamped HTML file.

### Web portal architecture and deployment

A lightweight dataset portal was implemented using a static web application using HTML, CSS and vanilla JavaScript. We configured the portal for deployment on Cloudflare Pages, while viewer assets were hosted separately on Cloudflare R2 using a public custom domain. Dataset metadata were stored in a JSON registry and loaded at runtime. Search and filtering are performed in-browser using title, description, citation and tag fields.

### Viewer externalization pipeline

Self-contained KaroSpace HTML exports were processed using a Python script to support large payloads. The script detects embedded JSON in blocks and JavaScript JSON assignment and then estimates the payload size and either then preserves it as a single file or externalizes the viewer into a directory format. In externalised mode, the data is written as chunked files with a generated manifest and loader runtime which then enables a reconstruction of JSON payloads when it comes time for the data to be loaded. Default settings use thresholds of 80MB with 50MB target chunk size.

### Cloud upload and thumbnail generation

Processed viewers were uploaded to Cloudflare R2 and public viewer links were then referenced in the portal metadata. Dataset preview images were generated from published viewer URLs using Playwright automation.

### Use of agentic coding tools

Development of KaroSpace and KaroSpaceBuilder was supported by agentic coding tools (Codex and Claude Code) for code drafting and debugging. All generated outputs were reviewed and validated by the authors.

## Data availability

The data used to generate the KaroSpace viewers are publicly available via the KaroSpace repository at https://github.com/christoffermattssonlangseth/KaroSpace. All data associated with this publication have been deposited in Zenodo under accession number 10.5281/zenodo.18670555. The interactive viewers can be accessed at https://karospace.se.

## Code availability

The source code required to generate KaroSpace viewers is available at https://github.com/christoffermattssonlangseth/KaroSpace and in KaroSpaceBuilder, https://github.com/christoffermattssonlangseth/KaroSpaceBuilder. The notebook used to integrate manually delineated polygons back into the AnnData object is available at: https://github.com/christoffermattssonlangseth/KaroSpace/blob/main/examples/add_polygons_back_to_anndata.ipynb.

## Acknowledgments

We acknowledge support from the National Genomics Infrastructure in Stockholm funded by Science for Life Laboratory, the Knut and Alice Wallenberg Foundation, and the Swedish Research Council. Part of the computation/data handling was enabled by resources provided by the National Academic Infrastructure for Supercomputing in Sweden (NAISS) and Swedish National Infrastructure for Computing (SNIC) at the Uppsala Multidisciplinary Center for Advanced Computational Science, partially funded by the Swedish Research Council through grant agreement no. 2022-06725 and no. 2018-05973. Part of the computing was also performed in the Linnarsson group Monod Linux cluster at MBB-KI, and we thank Peter Lönnerberg for maintenance and support. T.S. was supported by a Marie Skłodowska-Curie Actions post-doctoral fellowship. C.M.L. was supported by a Swedish Brain Foundation post-doctoral fellowship (PD2025-0508). Work in G.C.-B.’s research group was supported by the Swedish Research Council (grant 2019-01360 and Distinguished Professor grant 2023-00324), the European Union (Horizon 2020 Research and Innovation Programme/ European Research Council Advanced Grant SingleMS, grant agreement number 101096064), the Swedish Brain Foundation (FO2023-0032), the Swedish Cancer Society (Cancerfonden grant 23 2945 Pj 01 H), Knut and Alice Wallenberg Foundation (grant 2019-0089 and Wallenberg Scholar grant 2023-0280), the Göran Gustafsson Foundation for Research in Natural Sciences and Medicine, the Swedish Society for Medical Research (SSMF, grant JUB2019) and Karolinska Institutet.

